# CD45-mediated apoptosis and IL-2 receptor downregulation by serine proteases secreted from diarrheagenic bacteria

**DOI:** 10.1101/2025.03.20.644266

**Authors:** Yuexia Liao, Jorge L. Ayala-Lujan, Lixia Liu, Weijuan Gong, Guoqiang Zhu, James P. Nataro, Araceli E. Santiago, Fernando Ruiz-Perez

**Affiliations:** School of Nursing & Public Health, Yangzhou University, Yangzhou 225009, China; Academic Unit of Chemical Sciences, Autonomous University of Zacatecas, Zacatecas 98160, Mexico; School of Medicine, Yangzhou University, Yangzhou 225009, China. 11 Huaihai Rd, Guangling District, Yangzhou, Jiangsu, China, 225001; School of Veterinary Medicine, Yangzhou University, Yangzhou 225001, China; Child Health Research Center, Department of Pediatrics, University of Virginia School of Medicine, Charlottesville, VA 22908

## Abstract

Most enteropathogens secrete one or more members of the serine protease autotransporters of *Enterobacteriaceae* (SPATE). We previously demonstrated that SPATE cleaves various O-linked glycoproteins on leukocytes, including the tyrosine phosphatase CD45RO. SPATE impairs leukocyte functions and triggers apoptosis in activated T cells in vitro. Here, we show that SPATE produced by pathogenic *E. coli, Shigella*, and the mouse pathogen *Citrobacter rodentium* cleaves not only CD45RO but also CD45 isoforms containing exons A and B. We found that the cleavage of CD45 in primary T cells from both human and murine sources correlated with decreased IL2RA (CD25) surface expression in a concentration-dependent manner. SPATE did not cleave CD25 or affect T cell activation. However, SPATE requires CD45 expression for the depletion of CD25 in activated T cells, as SPATE did not significantly impact CD25 in the Jurkat J45.01 cell line, which lacks CD45. More importantly, we discovered that J45.01 cells resisted SPATE-mediated apoptosis, whereas apoptotic wild-type Jurkat cells exhibited decreased surface expression of CD25. Furthermore, we observed that mice infected with *C. rodentium* lacking SPATE displayed lower mortality, delayed intestinal colonization, reduced inflammatory cytokines, and decreased leukocyte infiltration in the lamina propria while having a higher number of CD25+ T cells compared to mice infected with wild-type CR or the CR SPATE mutant expressing Crc2 in trans. Our data suggest that SPATE-producing pathogens trigger T-cell apoptosis through CD45 via a mechanism akin to IL2 deprivation, demonstrating that SPATE can act as immunomodulators at various levels of the immune system.

**SIGNIFICANCE:** We have demonstrated for the first time that serine proteases (C2S) from clinically relevant pathogens, such as *E. coli* pathotypes and *Shigella*, can cleave leukocyte glycoproteins, including the tyrosine phosphatase CD45, which play crucial roles in cellular and immune functions. In this study, we discovered that C2S induces apoptosis in activated T cells through a previously unknown mechanism resembling IL-2 deprivation, mediated by CD45. Furthermore, we found that C2S is essential for bacterial virulence in vivo. This suggests that pathogens producing C2S may possess previously undescribed immunoregulatory functions that enhance their survival in the host and contribute to the disease process by eliminating T cells through the targeting of CD45 and the IL-2 receptor.

## Introduction

Enteric pathogens possess serine protease autotransporters of *Enterobacteriaceae* (SPATE), which resemble the trypsin-like superfamily of serine proteases and appear to have evolved to disrupt host homeostasis and alter the immune response as a mechanism of pathogenicity for survival in their environments^1^. They are produced by most, if not all, pathogens within the *Enterobacteriaceae* family. They have been shown to play crucial roles in bacterial pathogenesis^,1, 2^; however, the exact mechanisms of their involvement in immune evasion remain unclear and may result from SPATE cleavage of host cell receptors^3, 4^. We demonstrated that a subclass of SPATE, known as class-2 SPATEs (C2S), cleaves various leukocyte O-linked glycoproteins structurally similar to human mucin glycoproteins, including CD43, CD44, and CD45, which play diverse roles in cellular and immune functions ^4, 5^. More importantly, the cleavage of these glycoproteins impairs leukocyte functions and triggers T-cell apoptosis^4^. Pic is the archetypical C2S. We have demonstrated that Pic and several other C2S exhibit lectin activity, recognizing O-glycosylated Ser and Thr on glycoproteins^4, 5^. Upon binding to leukocytes, they induce neutrophil activation but impair chemotaxis and transmigration functions. Pic inhibits the neutrophil response to the chemokine IL-8 or fMLP (N-formyl-methionyl-leucyl-phenylalanine), as demonstrated in migration experiments, indicating impaired chemotaxis^4^. More intriguingly, Pic provoked apoptosis in activated T cells but not in resting lymphocytes, suggesting that Pic may impair T cell function during bacterial infections. Here, we further explore the relevance of C2S in bacterial pathogenesis and host immunomodulation by delving deeper into the phenomenon of cell death in T cells, identifying yet another mechanism of cell death induced by Pic, which relies on CD25 downregulation dependent on CD45 tyrosine phosphatase. CD25, also known as the α-subunit of the IL-2 receptor, regulates T cell function. Conversely, CD45 is an abundant and highly O-glycosylated protein found in various isoforms on the surface of all leukocytes, regulating innate and adaptive immune functions and programmed cell death in T and B-lymphocytes^6, 7^. Our data reveal another aspect of C2S as immunomodulators for CD45 and the IL-2 receptor in bacterial infections.

## RESULTS

### The serine protease Pic cleaves CD45 isoforms in T cells

We previously reported that the Pic serine protease degrades mucin-like O-glycosylated proteins on human leukocytes, including the protein tyrosine phosphatase CD45RO, which is also present in various isoforms in differentiated hematopoietic cells^4^. CD45 isoforms contain exons A, B, and C combinations or may lack exons (O)^6^. To further investigate the impact of Pic on CD45, we determined whether all CD45 isoforms were susceptible to Pic’s proteolytic activity. To do this, we treated approximately 1 million Peripheral Blood Mononuclear Cells (PBMCs) with either vehicle control or 1 to 0.1 µM of purified Pic. Following the treatment, PBMCs were analyzed by flow cytometry using antibodies specific to CD3, CD45RO, CD45RA, and CD45RB and with a pan-CD45 antibody, which recognizes all CD45 isoforms (Fig. 1A). We found that the extracellular domain of all CD45 isoforms containing A and B exons or O exons were removed from the surface of Pic-treated T cells (Fig. 1B). We confirmed that the pan-CD45 antibody recognized all CD45 isoforms regardless of cleavage status, suggesting that a common region of CD45 isoforms remains on the T cell surface. Our data indicate that Pic can remove all CD45 isoforms lacking or having combinations of A and B exons.

**Figure 1.**
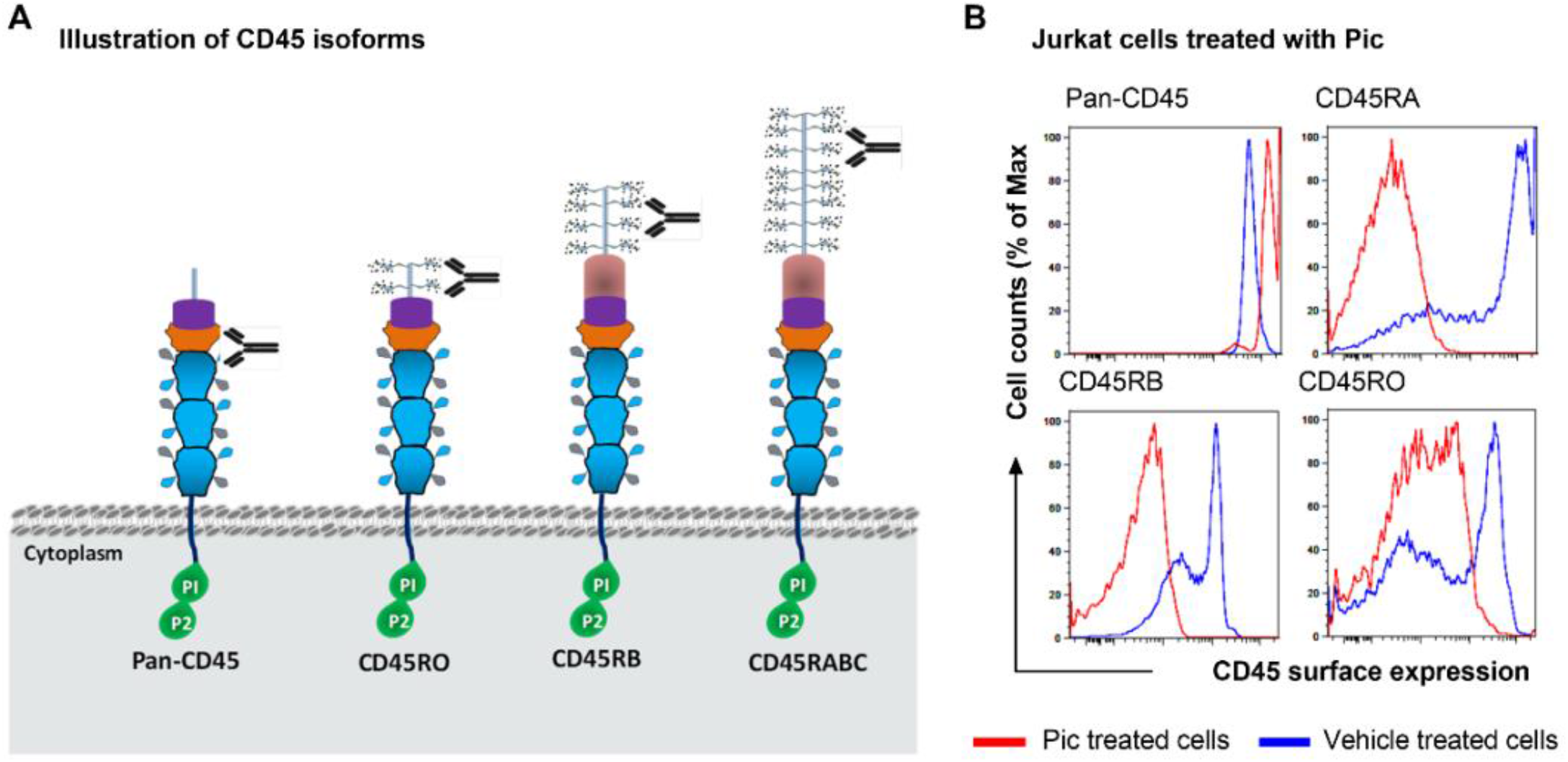
Pic cleaves all CD45 isoforms on T cells. **(A)** Illustration of CD45 isoforms and relative location of binding sites for antibodies used in this study. (**B**) A representative flow cytometry histogram of Jurkat cells treated with Pic for one h and analyzed for CD45 isoform surface expression using an anti-pan CD45 mAb and specific antibodies against each CD45 isoform.

### Pic serine protease affects T cells’ IL-2R (CD25) surface protein expression

We found that Pic’s proteolytic activity does not affect the viability of resting T cells but triggers apoptosis in a subset of activated T cells^4^. To study this phenomenon further, we activated the immortalized Jurkat T lymphocyte cell line overnight (∼18h) with a cell stimulation cocktail (PMA/Ionomycin) and recombinant IL-2 (Invitrogen). Subsequently, activated Jurkat cells were treated with increasing concentrations of purified Pic, a proteolytically inactive Pic protease (PicS258A)^8^, Crc2 protein (a Pic homolog produced by the mouse pathogen *C. rodentium*)^5, 9^, SepA (an unrelated SPATE protease)^10^, or only vehicle for 1 hour at 37°C. Next, the activation of Jurkat cells was assessed by analyzing the expression of early (CD69) and late (CD25) activation markers, along with the CD45O isoform using flow cytometry. We found that the PBS-treated Jurkat cells expressed 99% CD69, 65% CD25, and approximately 20% of the cells expressed CD45RO (Fig. 2A, B). To our surprise, we found that the degradation of CD45RO by Pic correlated with a decrease in CD25 surface expression in a Pic-concentration-dependent manner (Fig. 2A, B), but it did not affect CD69 expression (Fig. 2C). Activated Jurkat cells treated with Crc2 protein also exhibited degradation of CD45RO and decreased CD25 expression, whereas activated Jurkat cells treated with purified PicS258A or SepA did not show these effects (Fig. 2A, B). We found that the impact of Pic on CD25 was not dependent on how Jurkat cells were activated. Cells treated simultaneously with a stimulation cocktail and purified Pic exhibited the same effect on CD25 (Fig. 2D). We observed that pretreating Jurkat cells with Pic before activating them with the activation cocktail did not affect CD69 expression. However, it did affect the surface expression of CD25 (Fig. 2 E). This suggests that Pic does not influence T-cell activation but rather affects the expression of CD25. To investigate whether this effect also occurs in primary T cells and under a different activation method, human peripheral blood mononuclear cell (PBMC) T cells were activated through the T cell receptor (TCR) using monoclonal anti-CD3 and anti-CD28 antibodies for a co-stimulation signal. During this activation, the cells were treated with either purified SPATE or a vehicle control. We found that 70% of PBS-treated primary T cells expressed CD69, around 75% expressed CD25, and about 80% expressed CD45RO. Once again, we found that Pic and Crc2, but not PicS258 or SepA, degraded CD45RO in T cells in a concentration-dependent manner. This degradation correlated with decreased surface CD25 expression over time (Fig. 2G-H).

**Figure 2.**
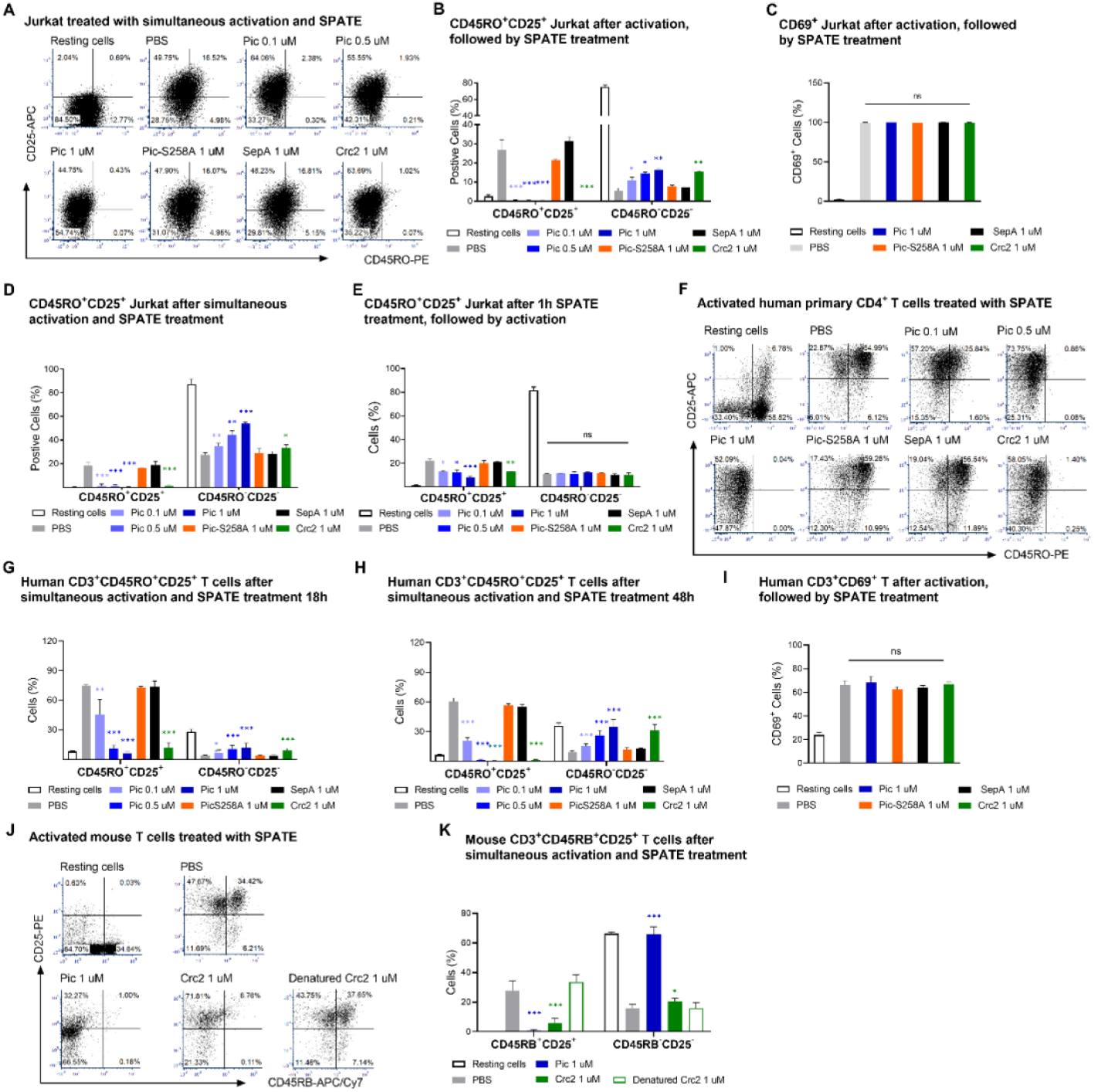
Pic and Crc2 serine proteases affect T cells’ IL-2Rα (CD25) surface protein expression. Jurkat cells, primary mouse T cells, and human T cells were activated with a cell stimulation cocktail and IL-2 or anti-human CD3 and CD28, respectively, before, simultaneously, or after SPATE (Pic, PicS258A, SepA, Crc2) treatment. A representative flow cytometry histogram for CD45RO+CD25+ Jurkat cells after simultaneous activation and SPATE treatment is shown in (A). Statistical analysis of CD69+, CD45RO+CD25+, and CD45RO-CD25-Jurkat cell populations following cell activation and SPATE treatment is displayed in (**B, C**), simultaneous cell activation with SPATE treatment in (**D**), or one hour of SPATE treatment before cell activation in (**E**). A flow cytometry histogram (**F**) and statistical analysis (**G, H, I**) of human CD69+, CD3+CD45RO+CD25+ and CD3+CD45RO-CD25-cell populations after 18 hours (**H**) or 48 hours (**I**) of cell activation and SPATE treatment are presented. Figures **J** and **K** illustrate a representative flow cytometry histogram and statistical analysis of mouse splenic CD45RB+CD25+ and CD45RB-CD25-T cells that underwent simultaneous cell activation and SPATE treatment. The data represent three independent experiments, with values expressed as mean ± SD and analyzed against activated T cells treated only with vehicle (PBS) using an unpaired t-test. *P < 0.05, **P < 0.01, and ***P < 0.001. ns: not significant.

Lastly, we investigated the effects of Pic and Crc2 on CD45 and CD25 surface expression in activated T-cells derived from murine spleen. This experiment used a monoclonal antibody that recognizes CD45 isoforms containing the B exon. As expected, activated mouse T-cells treated with PBS or heat-denatured Crc2 expressed CD45RB isoforms. In contrast, only 1 to 7 % of T-cells treated with Pic or Crc2 showed CD45B surface expression (Fig. 2 J, K). Notably, the degradation of CD45RB correlated with reduced CD25 surface expression (Fig. 2K).

### The depletion of CD25 on T cells is not due to the proteolytic activity of SPATE

To determine whether the loss of CD25 surface expression resulted from the degradation of CD25 by Pic or Crc2, we measured CD25 levels in Jurkat cells after SPATE treatment. For this purpose, Jurkat cells were activated and treated with 1 µM Pic, PicS258A, SepA, Crc2, or vehicle only for 18 hours at 37°C as previously described. After the activation period, we quantified the levels of CD25 in Jurkat cell lysates and supernatants using ELISA, SDS-PAGE, and western blot. We did not observe any significant differences in CD25 protein levels in the supernatants of Jurkat cells in the ELISA assay (Fig. 3A). Additionally, we did not detect any degraded products of CD25 in either wild-type Jurkat or J45.01cell lines ^11^, as determined by western blot analysis (Fig. 3B). Furthermore, recombinant CD25 was not degraded by purified Pic or other SPATE, as shown by SDS-PAGE (Fig. 3C).

**Figure 3.**
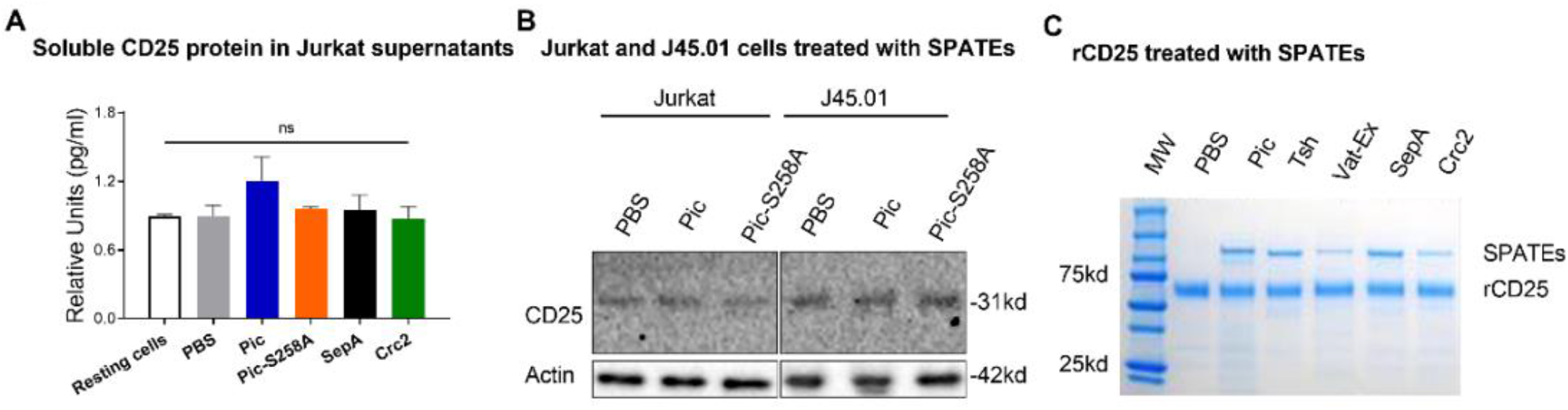
The decrease of CD25 on the T cell surface is not due to Pic-mediated CD25 cleavage. Activated Jurkat cells were treated with SPATE for 18 h, as previously. Following the treatment, the amount of soluble CD25 protein in Jurkat cells’ supernatants was measured using an ELISA assay (**A**) and in cell lysates by western blot using an antibody against CD25 and β-actin as a loading control (**B**). CD25 protein expression was also analyzed in the Jurkat cell line J45.01 (**B**, J45.01). The possibility of CD25 cleavage by SPATEs was also assessed by incubating ∼1ug of recombinant CD25 with 1-2 μM of SPATEs at 37°C overnight, followed by SDS-PAGE analysis (**C**). Lastly, CD25 gene expression was evaluated by qRT-PCR (**D**).

### The reduction of CD25 on the surface of T-cells caused by SPATE depends on CD45

Our data suggest that reduced CD25 expression in Pic-treated T cells is associated with CD45 degradation. To determine the direct role of CD45 in this process, we used the CD45-defective Jurkat J45.01 cell line, which expresses less than 10% of CD45 compared to the parental Jurkat cell line, resulting in defective signal transduction^11^. We initially examined the presence of CD45RO in both parental and J45.01 cells under resting and activation conditions. Our results confirmed minimal expression of CD45RO in J45.01 cells, both before and after T cell activation (Fig. 4A). In contrast to J45.01, wild-type Jurkat cells exhibited high levels of CD45RO expression prior to activation, with even higher levels observed after activation (Fig. 4A). Next, we treated activated J45.01 cells with SPATEs or only vehicle and analyzed activation markers by flow cytometry (Fig. 4B-D). We observed no significant differences in the CD45RO+CD25+ J45.01 cell population following Pic treatment compared to the other treatments (Fig. 4B, C). When comparing the percentage of CD25+ cells in wild-type Jurkat and J45.01 cells treated with increasing concentrations of Pic, we noted a significant reduction of CD25 in the parental Jurkat strain compared to J45.01 (Fig. 4D). Only at high concentrations of Pic was there approximately a 10% reduction in CD25 expression in J45.01 cells, which may correspond with the residual expression of CD45 in these cells. Cell surface CD69 expression also remained unaffected in activated J45.01 (Fig. 4E).

**Figure 4.**
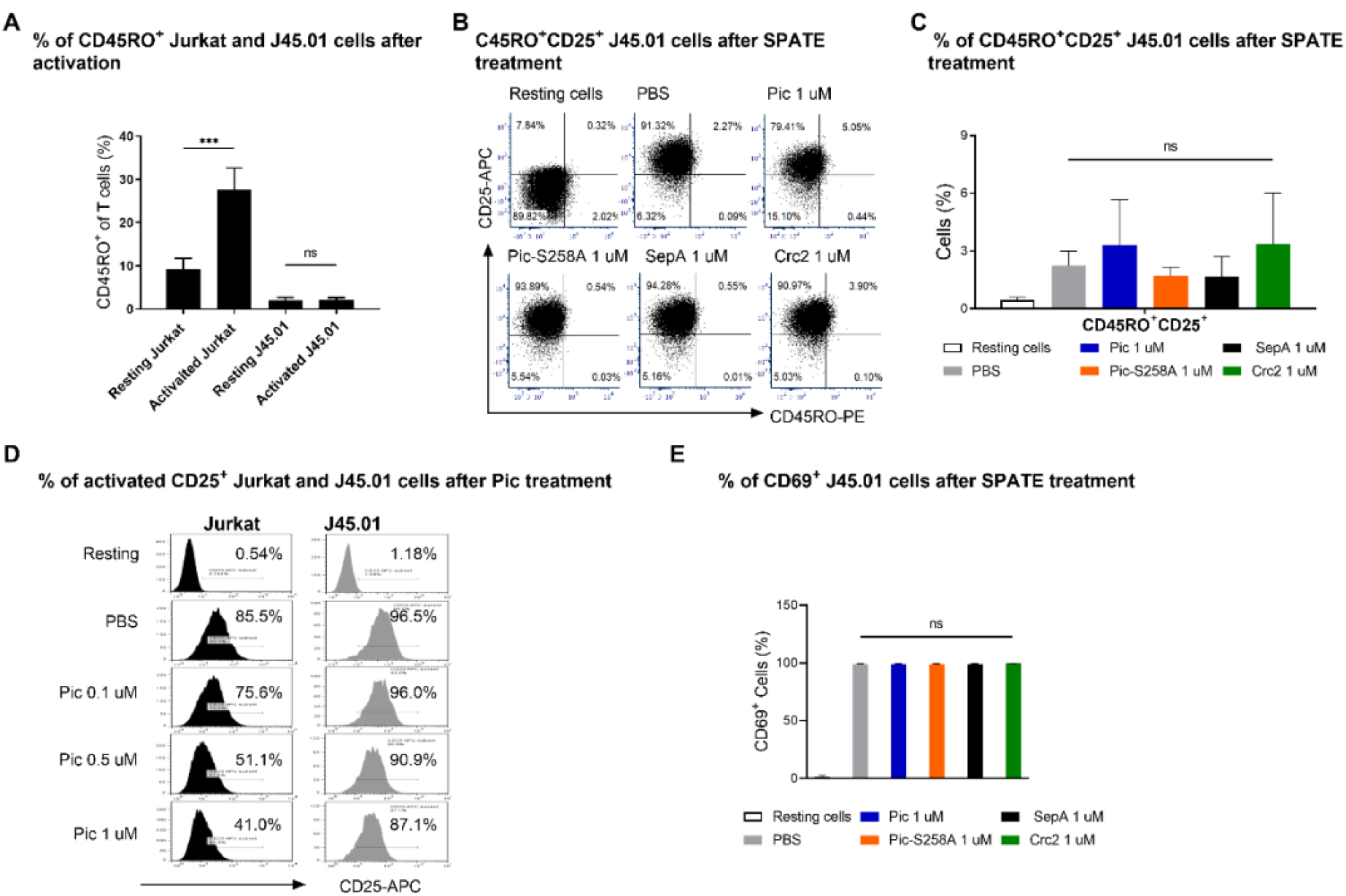
The CD25 surface reduction in T-cells due to Pic depends on CD45. CD45-deficient (J45.01) Jurkat cells were activated with a Cell stimulation cocktail and IL-2 and treated with SPATE (Pic, PicS258A, SepA, Crc2) for 18h. (**A**) Comparison of CD45 expression in resting and activated Jurkat and J45.01 cells. (**B**) A representative flow cytometry histogram for CD45RO^+^CD25^+^ J45.01 cells after SPATE treatment. (**C, E**) Statistical analysis of CD69^+^ and CD45RO^+^CD25^+^ J45.01 cell population after SPATE treatment. (**D**) Percentage of CDCD25+ Jurkat and J45.01 cells after treatment with various concentrations of Pic. Data represent three independent experiments, and values are expressed in mean ± SD and analyzed by unpaired t-test. \**P* < 0.05, \*\**P* < 0.01, and \*\*\**P* < 0.001. ns: not significant.

### Inhibition of CD45 phosphatase activity causes CD25 downregulation in activated T cells

CD45 tyrosine phosphatase is central to immune cell activation and cytokine receptor signaling ^6, 12^. To determine whether inhibiting CD45 phosphatase activity with a CD45 chemical inhibitor produces the same effect on CD25 as treatment with Pic, we treated Jurkat cells with the CD45 inhibitor both in the absence and presence of Pic and measured CD25 surface expression via flow cytometry. We found that inhibiting CD45 led to a decrease in CD25 surface expression in Jurkat cells, and Pic acted as an agonist of CD25 suppression when added alongside the CD45 inhibitor (Supplemental Fig.1). This suggests that the reduction in CD25 surface expression on activated cells treated with Pic may be due to CD45 inactivation.

### Pic causes T-cell apoptosis dependent on CD45

We previously reported that pretreatment of T cells with Pic causes apoptosis in activated but not in resting cells ^4^. To investigate whether T cell apoptosis induced by Pic results from CD45 cleavage or CD25 surface depletion, activated Jurkat and J45.01 cells were treated with SPATEs or a control vehicle, as previously described. Following treatment, the cells were stained with Annexin V and propidium iodide (PI) to assess apoptosis and analyze the expression levels of CD45 and CD25. As anticipated, we observed that Pic and Crc2 triggered apoptosis in Jurkat T cells in a concentration-dependent manner. This correlated with an increase in Annexin V/PI staining and a decrease in CD25+ surface expression (Fig. 5A, B, and G). The confocal analysis also revealed many apoptotic nuclei on Pic-treated Jurkat cells, correlating with a reduced CD25 expression (Fig. 5C, D). Unexpectedly, we found that neither Pic nor Crc2 caused significant apoptosis in J45.01, nor did they considerably affect the expression of CD25 in Annexin+ cells (Fig 5E, F, and G). Our data suggest that apoptosis triggered by Pic in activated T cells is dependent on CD45, potentially due to the downregulation of CD25, similar to the apoptosis caused by IL-2 deprivation^13-17^.

**Figure 5.**
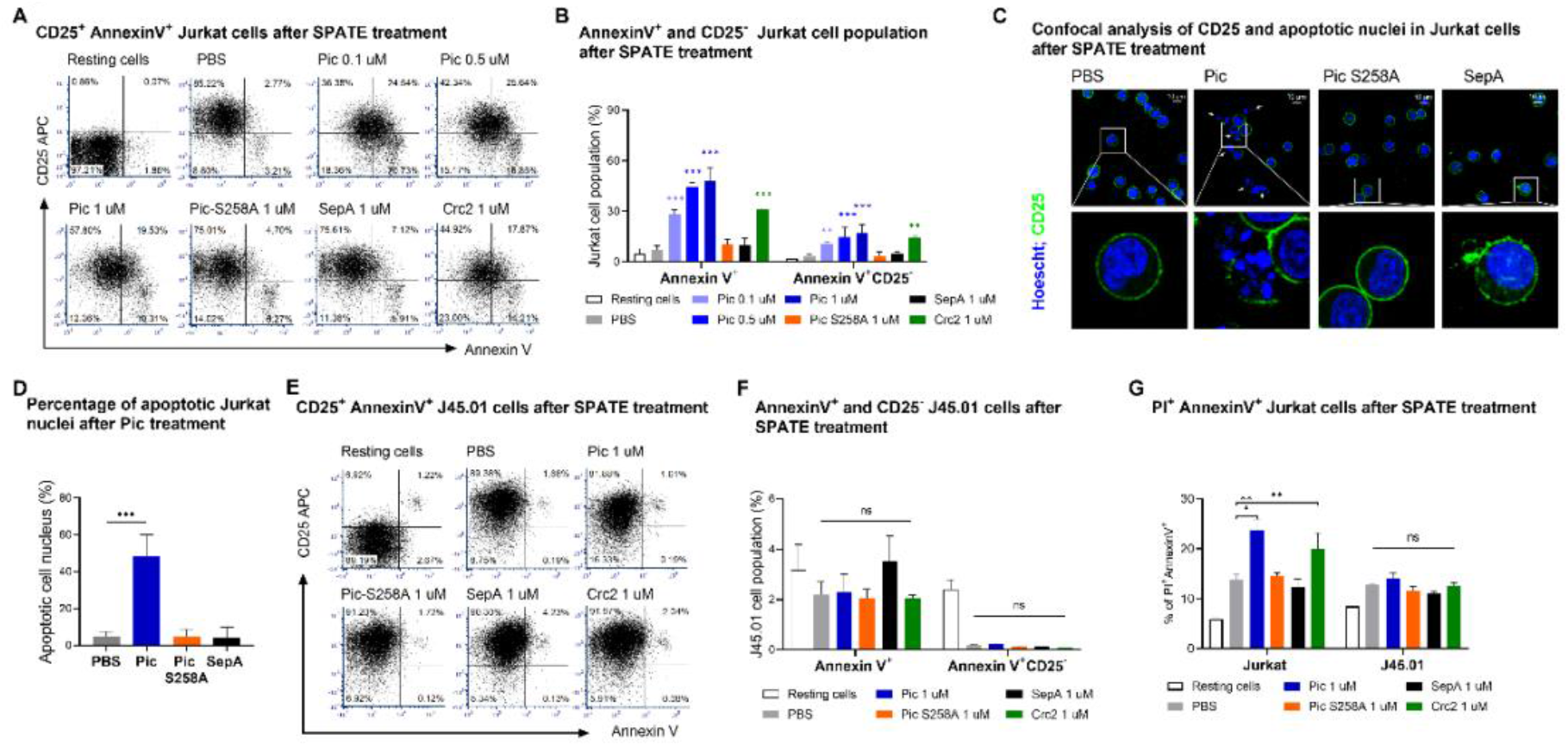
Pic and Crc2 cause T-cell apoptosis in Jurkat T cells but not in J45.01 cells. Jurkat and J45.01 cells were activated with a cell stimulation cocktail and IL-2 and treated with SPATE (Pic, PicS258A, SepA, Crc2) for 18h. Representative flow cytometry histograms for AnnexinV^+^ CD25^+^ (PI-negatively gated) Jurkat (**A**) and J45.01 (**E**) T-cells after SPATE treatment are shown. Statistically significant differences in AnnexinV^+^ and CD25^-^ Jurkat (**B**) and J45.01 (**F**) cells after cell activation and SPATE treatment are shown. Statistically significant differences in AnnexinV^+^/PI^+^ Jurkat and J45.01 T cells after cell activation and SPATE treatment are shown in **G**. Percentage of apoptotic Jurkat cell nuclei after SPATE treatment was visualized and quantified by confocal microscopy by staining with Hoechst (DNA) and with an Alexa-conjugated anti-CD25 mAb (**C, D)**. Data represent three independent experiments, and values are expressed in mean ± SD and analyzed by unpaired t-test. \**P* < 0.05, \*\**P* < 0.01, and \*\*\**P* < 0.001. ns: not significant.

### Crc2 plays a role in *C. rodentium* pathogenesis and inflammatory response

We sought to determine the relevance of SPATE in the immune response during infection. We, therefore, employ *C. rodentium* (CR), which is widely used as a surrogate model of human pathogens to investigate innate and adaptive mucosal immune responses to bacterial intestinal infection^18^. We previously identified three SPATEs in CR, which are Crc1, Crc2, and AdcA ^5, 9^. We and other groups have observed that deleting a single SPATE in CR renders similar hyper virulent phenotypes and a higher inflammatory response despite having different substrate repertoire ^9, 19 5^. To study the role of a single SPATE in CR infection, we generated CR isogenic strains lacking all three SPATEs and complementing with a low copy plasmid expressing only Crc2, the closest Pic homolog^5^. Accordingly, groups of mice were mock infected or infected with wild-type CR, CR lacking SPATEs (CR Δ_SPATE_), or CR Δ_SPATE_ complemented with a low copy plasmid expressing Crc2 (pCr2). Surprisingly, we found that mice infected with a CR lacking SPATEs (CR Δ_SPATE_ mutant) showed reduced signs of diseases, including less mouse mortality (Fig. 6A), weight loss (Fig. 6B), crypt hyperplasia (Fig 6D, E), and inflammatory cytokines (Fig. 6F-H) than mice infected with the wild-type CR strain or the mutant expressing only Crc2. The CR strains that lacked SPATE exhibited reduced intestinal colonization during the early stages of infection (Fig. 6C, Day 3). All CR strains showed comparable colonic bacterial burdens by the mid-infection stages (Days 6-8). However, the CR_ΔSPATE_ mutant began to show clearance signs by day 10. Moreover, animals infected with the CR_ΔSPATE_ mutant showed significantly reduced neutrophil and macrophage infiltrates (Fig. 6I, J) but higher T cells in the lamina propria (LP)(Fig 6. K-M) than mice infected with the wild-type or the Cr2-only strain at days 7 and 11 post-infection. The levels of pro-inflammatory cytokine response between the mouse groups correlated with leukocyte infiltrates. In contrast, at late infection stages, the high number of T cells in CR_ΔSPATE_ correlated with higher levels of IFN-gamma (Fig. 6H).

**Figure 6.**
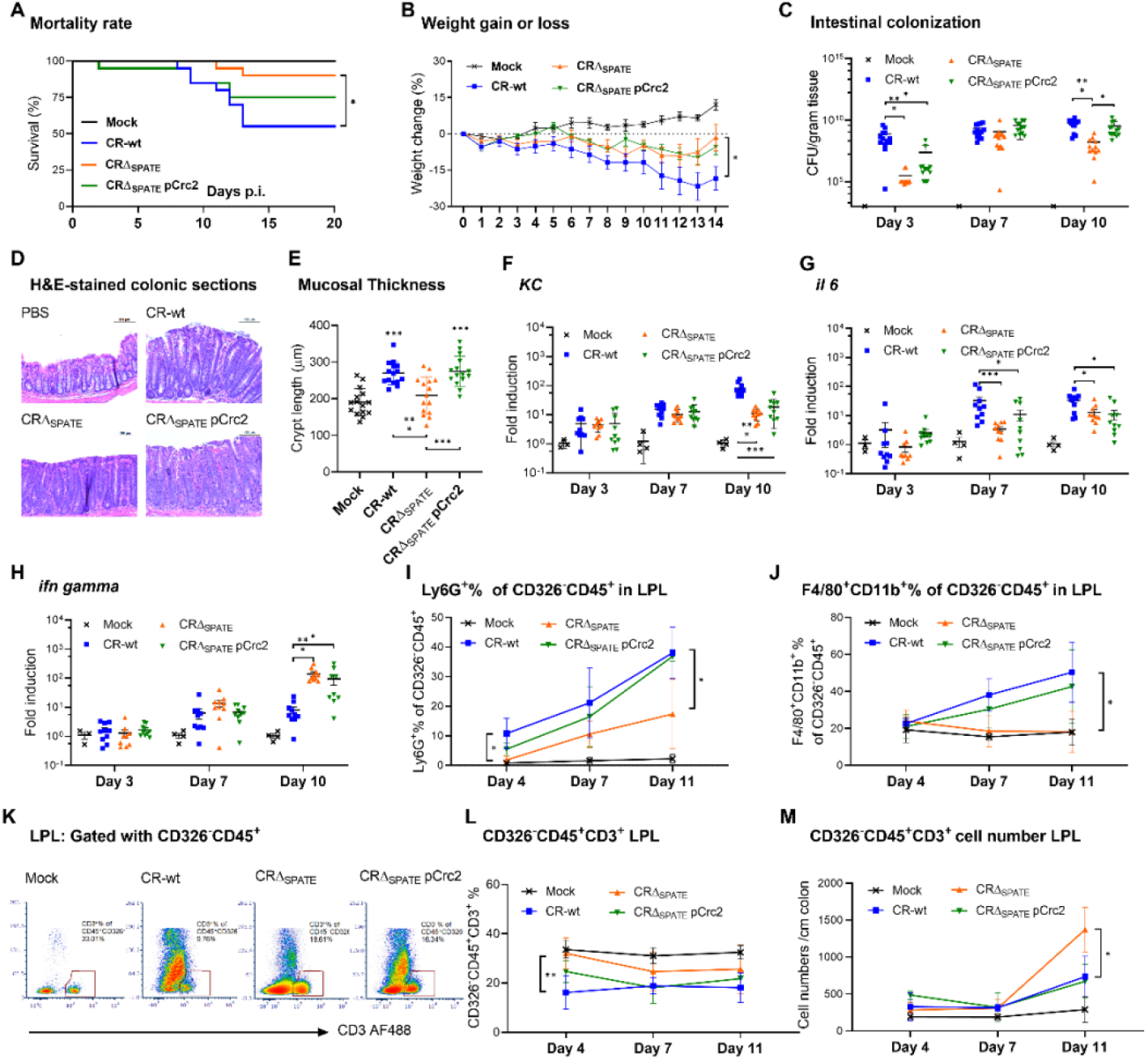
Crc2 influences C. rodentium pathogenesis and the host inflammatory response. Groups of C57BL/6 mice were mock-infected or infected with 10E10 CFU of wild-type Citrobacter rodentium (CR-wt), its isogenic mutant lacking SPATEs (CRSPATE), or the mutant complemented with the pCrc2 plasmid encoding Crc2 (CRSPATE pCrc2). Mice were monitored for survival (A), weight gain or loss (B), bacterial CFUs in the distal colon (C), crypt hyperplasia as indicated by H&E-stained colonic sections and crypt length measurements (D, E), Lamina propria (LP) cytokines (F, G, H), neutrophil and macrophage infiltrates (I, J), and the percentage or cell number of T-cell infiltrates (K-M). LP leukocytes were isolated at day 11 PI and analyzed using Flow Cytometry, normalizing to 10,000 counting beads and gating on live cells. T cells, PMNs, and macrophages are gated on CD326-/CD45+/CD3+, CD326-/CD45+/Gr1+, and CD326-/CD45+/CD11b+/F4/80+ cells, respectively. Survival was analyzed with the log-rank (Mantel-Cox) test. Weight loss, colonization, cytokine expression, and leukocyte infiltrates were analyzed using two-way ANOVA with multiple comparisons or an unpaired Student’s t-test. Asterisks denote statistically significant differences between the groups: *P < 0.05, **P < 0.01, and ***P < 0.001. ns: not significant.

### Reduced CD45RB+ and CD25+ T cells in mice infected with *C. rodentium* expressing SPATE

Since Crc2 degraded CD45 isoforms and affected the expression of CD25 in vitro, we sought to determine if these markers are also affected in LP T cells during CR infection. Therefore, we infected mice with wild-type CR, CR Δ_SPATE_, or CR Δ_SPATE_ pCr2 and analyzed the CD45RB+ and CD25+ T-cell subpopulations at days 4, 7, and 11 post-infection. We did not observe significant differences in CD44+, CD162+, or CD45RA+ (data not shown) in T cell populations between the mouse groups. Still, significantly higher CD45RB+ and CD25+ T cells were observed in mice infected with the CR Δ_SPATE_ strain (Fig. 7A-C), suggesting a direct or indirect effect of SPATE, including Crc2, on these T cell populations.

**Figure 7.**
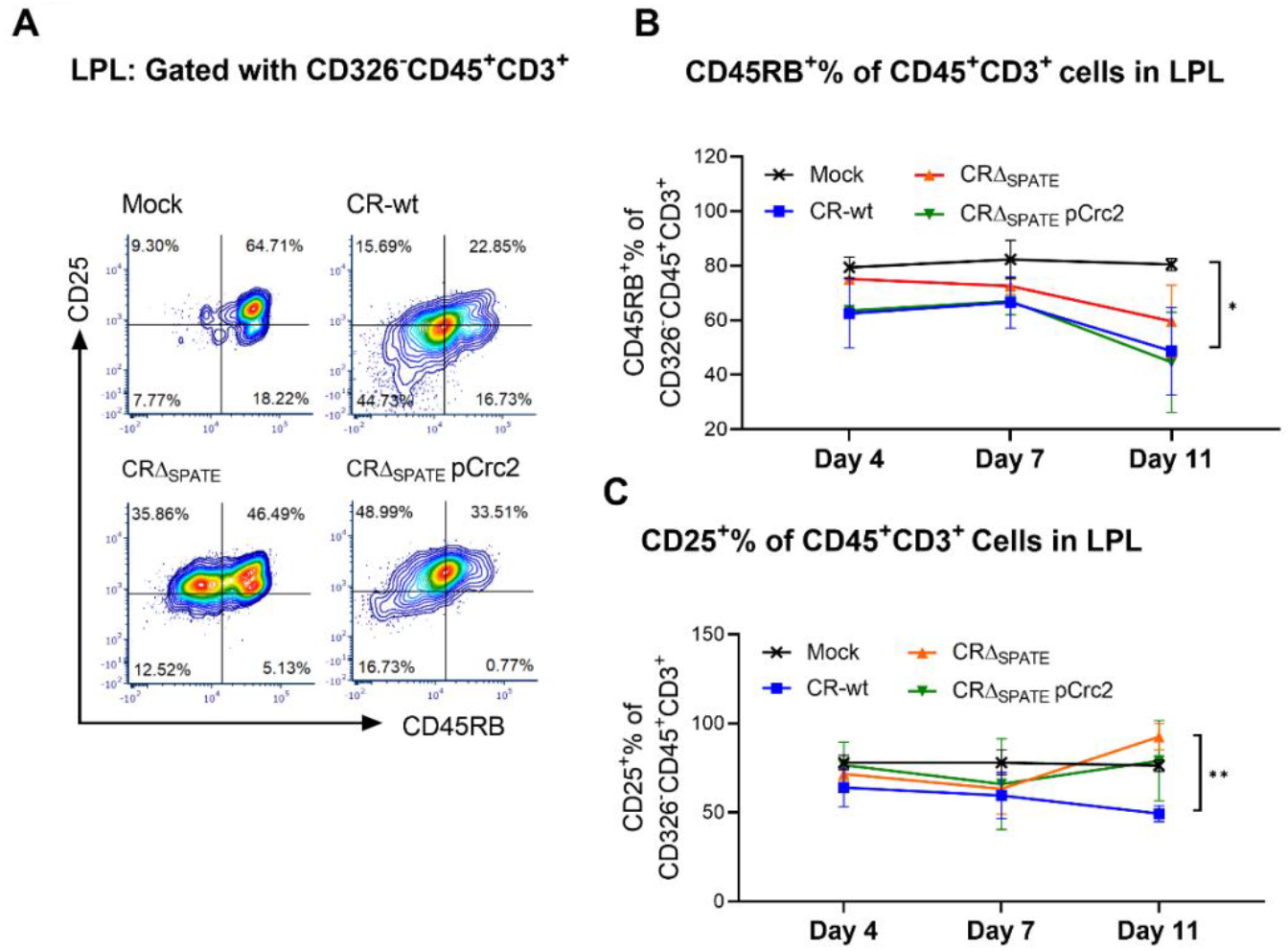
Reduced CD45RB^+^ and CD25^+^ populations in colonic Lamina Propria of mice infected with wild-type CR compared to CR Δ_SPATE_ mutant. Groups of 10 C57BL/6 mice were mock-infected or infected with CR derivatives as in Fig. 6. Lamina propria leukocytes were isolated on Day 4^th^, 7^th^, and 11^th^ and subjected to flow cytometry analysis. A representative FC histogram of the T-cell population gated on CD326^-^CD45^+^CD3^+^ from live cells is shown in **A**. CD326^-^CD45^+^CD3^+^ T cell subpopulations were further gated on CD45RB+ (**B**) and CD25+ (**C**). Data are represented as mean ± SD and analyzed with an unpaired t-test. \**P* < 0.05, \*\**P* < 0.01, and \*\*\**P* < 0.001. ns: not significant).

## Discussion

We have previously demonstrated that the prototype SPATE, Pic, cleaves many mucin-like O-linked glycoproteins on human leukocytes, including the tyrosine phosphatase CD45RO. In vitro, Pic induces apoptosis in activated T cells but not in resting lymphocytes. Given the prevalence of CD45 in leukocytes and its significance in T cell function, we further explored how CD45 targeting by Pic influences T cell activation and viability. We also investigated the effects of the Pic homolog Crc2 from *C. rodentium* in the context of infection.

CD45 is a highly glycosylated tyrosine phosphatase receptor that exists in multiple isoforms, such as CD45RA and CD45RB in naïve T cells, CD45RO in memory T cells, and CD45RABC in B cells ^20, 21^. The glycosylation state of CD45 influences the recognition by various binding partners ^22-24^. It influences intracellular signaling via the cytoplasmic tyrosine phosphatase domain, positively or negatively affecting the T cell’s response to an antigen^23, 24^. We found that Pic cleaves not only CD45RO but also the O-glycosylated extracellular domain of CD45 isoforms containing exons A and B. Pic’s cleavage of the extracellular domain of various CD45 isoforms was efficient, completely removing it from the T cell surface at all tested Pic concentrations. Additionally, we observed that the CD45 pan-antibody, which does not recognize the O-glycosylated exons, still identified CD45 even after Pic treatment, indicating that a truncated CD45 extracellular domain remains on the T cell surface.

To investigate the impact of CD45 cleavage by Pic on T-cell activation and apoptosis, we conducted experiments in which Jurkat T cells and human and murine primary T cells were incubated with LPS-free purified Pic before, simultaneously, or after T-cell activation. In addition to the cleavage of CD45, we observed that the early activation marker CD69 was overexpressed on the T cell surface, as anticipated. However, the late activation marker CD25 was depleted from the T cell surface in a Pic concentration-dependent manner. Depletion of CD25 by Pic was observed under all tested conditions, whether before or after T cell activation, in both immortalized and primary cells derived from mice and humans. This effect did not depend on the strategy used to activate the cells, as the PMA/ionomycin activation cocktail, which bypasses the T cell receptor (TCR), or the anti-CD23-CD28 cocktail, which partially mimics stimulation by antigen-presenting cells, yielded the same results on CD25 depletion. Since Pic has been shown to cleave other glycosylated proteins, we reasoned that Pic could have cleaved CD25. However, ELISA and western blot experiments searching for cleaved forms of CD25 in T cell supernatants or cell lysates did not indicate that CD25 was cleaved. We also did not observe Pic cleaving human recombinant CD25.

To investigate the correlation between CD45 cleavage and CD25 downregulation by Pic, we utilized the Jurkat cell line J45.01, deficient in CD45 expression. Surprisingly, we discovered that Pic was unable to deplete CD25 in the J45.01 cell line effectively as in the parental Jurkat cell line; a high concentration of Pic only depleted about 10% of CD25 in J45.01, compared to 50% in the parental Jurkat cell line in the period analyzed. The low percentage of J45.01 cells showing depletion of CD25 may be due to this cell line retaining 5-8% CD45 expression relative to the parental Jurkat cell line^24^. This finding suggests that CD25 downregulation is linked to the degradation of CD45 by Pic. This led us to test whether inhibiting CD45 signaling in activated T cells with a CD45 inhibitor could induce CD25 depletion. In fact, approximately 2.5 µM of CD45 inhibitor (Sigma) resulted in significant CD25 depletion in activated Jurkat cells, and co-treatment with Pic had an even more substantial effect on CD25 depletion.

CD45 has been shown to modulate T-cell apoptosis. T lymphocytes expressing single CD45RABC or CD45RO isoforms exhibit increased apoptosis, and the degree of apoptosis is inversely correlated with CD45 expression ^25^. We, therefore, sought to determine whether Pic causes T-cell apoptosis by interacting with CD45. Surprisingly, when treated with Pic, the CD45-deficient J45.01 cell line did not undergo significant apoptosis as the parental Jurkat cell line. Other studies have shown that ligation of the extracellular portion of CD45 (RO, RA, or RB) using monoclonal antibodies can trigger apoptosis in T cells^7^. Similarly, the binding of CD45 with Galectin-1 and Galectin-3, both members of a conserved lectin family associated with immune regulation, triggers the death of human and murine thymocytes and activated human T cells ^26,27^. Indeed, T cells deficient for CD45 expression, like the J45.01 cell line, were resistant to galectin-3-induced T cell death, and the modulation of apoptosis by galectin-1 depended on O-glycosylation ^26^. Our data suggest that Pic and possibly other class-2 SPATE, such as Crc2, induce apoptosis in T cells by interacting with and degrading the O-glycosylated CD45 receptor. This interaction leads to CD25 depletion in activated T cells, which may induce cell apoptosis via a mechanism similar to IL-2 deprivation ^13, 14, 16, 17^. In vitro studies have shown that IL-2 is essential for regulating T-cell antigen response. CD25, the IL-2 receptor alpha (IL-2Rα), is part of the interleukin-2 (IL-2) receptor complex, along with interleukin-2 receptor beta (IL-2Rβ) and interleukin-2 receptor gamma (IL-2Rγ). All three components are necessary for the receptor complex to function properly. Activated T cells that cannot produce or respond to IL-2 due to targeted mutations in the IL-2 or IL-2R genes undergo programmed cell death ^14, 17^.

To examine the immunomodulatory role of SPATE in the context of infection, we used the *C. rodentium* (CR) mouse pathogen as a genuine infection model, widely used as a surrogate for human pathogens to investigate innate and adaptive mucosal immune responses to bacterial infection ^18^. We have characterized three SPATE proteases in CR, termed Crc1, Crc2, and AdcA ^5, 9^. Like other class-2 SPATE they have unique or overlapping substrates. Therefore, to investigate the individual roles of SPATE, we deleted all three SPATE genes from the CR strain, and the resulting triple mutant was complemented with a low-copy plasmid expressing Crc2, the closest Pic homolog. We found that mice infected with wild-type CR or strain-producing only Crc2 succumbed more readily to infection and presented a higher bacterial burden, neutrophil and macrophage infiltrates, and proinflammatory cytokine in the lamina propria. However, mice infected with a CR expressing no SPATE presented a higher T cell population, interferon-γ, CD45B+ cells, and significantly more CD25+ positive cells in the lamina propria at day 11 of infection. Overall, our data indicate that SPATE-producing pathogens have developed previously unrecognized key immunoregulatory functions that enhance their survival in the host and contribute to pathogenesis (Fig. 8). Advancements in understanding the molecular mechanisms of SPATEs’ immunomodulatory roles may also illuminate the emerging functions of mammalian lectins and shedases in immunoregulation.

**Figure 8.**
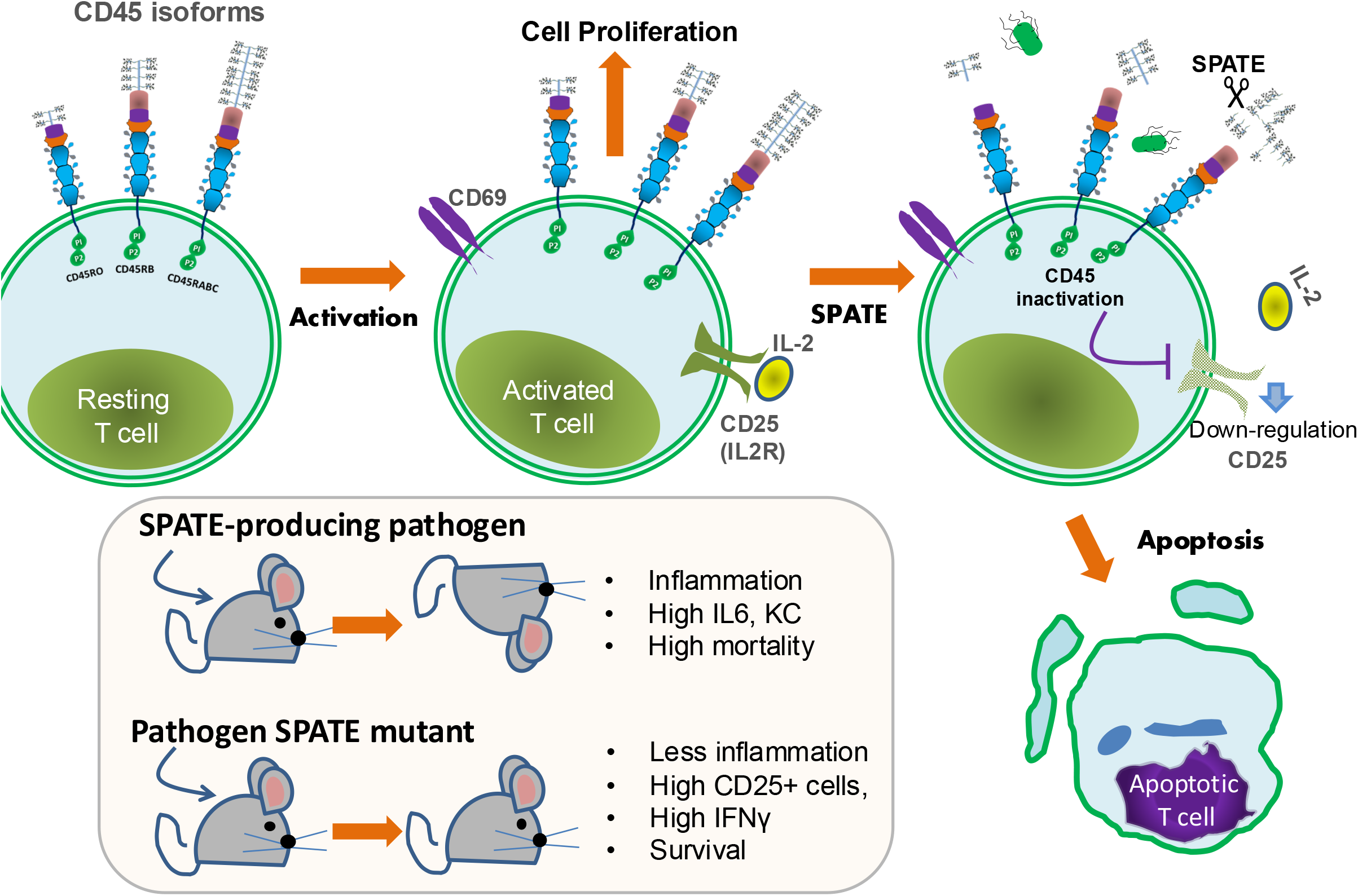
CD45-mediated apoptosis and IL-2 receptor downregulation by C2S. Serine proteases (C2S) from clinically relevant pathogens can cleave leukocyte glycoproteins, including CD45. In this study, we discovered that C2S induces apoptosis in activated T cells through a previously unknown mechanism that resembles IL-2 deprivation mediated by CD45. Furthermore, we found that C2S is essential for bacterial virulence in vivo. This suggests that pathogens producing C2S may have evolved previously undescribed immunoregulatory functions that enhance their survival in the host and contribute to the disease process

## Material and Methods

### Cell lines and human primary cells

The Jurkat T cell line (Clone E6-1, ATCC TIB-152) and J45.01 (derivative from E6-1)^11^ were typically cultured in RPMI 1640 medium supplemented with 10% heat-inactivated fetal bovine serum (FBS) and antibiotics: 100 U/ml penicillin and 100 µg/ml streptomycin, and incubated at 37°C in a moisture-rich environment of 95% O2 and 5% CO2. Fresh heparinized blood from healthy human donors, obtained from All Cells LLC (Alameda, CA, 94502), was mixed in a 1:1 ratio with dextran/0.9% NaCl and allowed to stand undisturbed for 30 minutes. To remove any residual erythrocytes, the cell fraction from the leukocyte-rich plasma was centrifuged at 1500 rpm for 15 minutes and then resuspended in 12 ml of cold, sterile water for 25 seconds, followed by the addition of 4 ml of 3% NaCl. Red blood cell debris was eliminated by centrifugation at 1500 rpm for 10 minutes, followed by a wash with HBSS buffer lacking Ca++/Mg++ (Invitrogen).

### Activation of Jurkat and primary T cells

Jurkat cells were activated with 1X Cell Stimulation Cocktail (eBioscience) and 5 ng/ml of IL-2 for 18 hours at 37°C in a humid environment of 95% O2 and 5% CO2. Mouse splenic T cells and human PBMCs were activated using 5 ng/ml of IL-2 and 0.5 ug/ml of anti-CD3 and anti-CD28 monoclonal antibodies for up to five days at 37°C in an environment of 95% O2 and 5% CO2.

### CD25 Detection in Jurkat Cell Supernatants

After treatment with SPATEs, the concentration of soluble CD25 protein in the supernatants of Jurkat cells was measured in triplicates using the Human CD25/IL-2R alpha Quantikine ELISA Kit (R&D Systems), according to the manufacturer’s recommendations.

### Purification of SPATE Proteins

The expression and purification of Pic, Pic-S258A, SepA, and other SPATE proteins from minimal clones in *E. coli* HB101 have been detailed previously^4^. Anion-exchange purified proteins were treated with the EndoTrap® HD Endotoxin Removal System from BioVendor to remove LPS according to the manufacturer’s instructions, dialyzed in PBS at pH 7.0, and stored in small aliquots at -80°C.

### Construction of a *C. rodentium* SPATE mutant

The *C. rodentium* SPATE mutant was created by sequentially deleting the genes encoding crc1, crc2, and adcA using the λ Red recombinase system. PCR fragments containing a kanamycin resistance gene were generated, with 200 bp nucleotide sequences flanking the regions upstream and downstream of the crc1, crc2, and adcA genes (supplemental table-1). These fragments were then electroporated into the *C. rodentium* strain ATCC 51459, which expresses the λ Red recombinase system through the plasmid pKM200. Mutants were selected on LB agar plates containing kanamycin (50 mg/ml). All gene deletions were verified by PCR amplification using primers located outside the disrupted gene regions. The crc1/crc2/adcA triple mutant was complemented with the pACYC184-Crc2 plasmid.

### Infection of mice and histology

The *C. rodentium* model was utilized as described^9^. Briefly, 3- to 4-week-old C57BL/6 mice were obtained from Jackson Laboratories (Bar Harbor, ME). Ten mice per group were inoculated with approximately 10^10 CFU of wild-type *C. rodentium*, the SPATE (*Δcrc1, crc2, adcA*) mutant, or the complemented SPATE mutant strain bearing the pACYC184-Crc2 plasmid, all suspended in 200 μl of phosphate-buffered saline (PBS). The organisms were administered via oral gavage using a 20-gauge intubation needle. Control animals received 200 μl of sterile PBS. At specified time points after inoculation, 3 cm segments of the distal colon were collected aseptically, gently rinsed with PBS, weighed, and homogenized in PBS. The number of viable bacteria per gram of tissue was determined through serial dilutions of the samples on selective antibiotic-based media.

Histology: Colonic tissues were fixed in 4% paraformaldehyde and processed for wax embedding. Cross-sections of the colon (5 μm) were sliced, mounted on slides, and stained with hematoxylin and eosin (H&E).

### Ethics statement

Animal experiments were performed in accordance with the Guide for the Care and Use of Laboratory Animals of the National Institutes of Health and with the permission of the American Association for the Assessment and Accreditation of Laboratory Animal Care. The protocol was reviewed and approved by the Institutional Animal Care and Use Committee of the University of Virginia (protocol number 3894).

### Analysis of Leukocyte Populations in the Distal Colon

The leukocyte infiltrates in the colon lamina propria were analyzed as previously described^9^. Briefly, on days 1, 3, 5, and either 10 or 11 after inoculation, mice were euthanized, and a 3 cm segment of the distal colon was excised and placed in a petri dish containing Hanks’ Balanced Salt Solution (HBSS). The mesentery, fat, and feces were carefully removed from the distal colon, which was then opened lengthwise and cut into 0.5 cm segments. These segments were transferred to 20 ml of pre-digestion buffer (HBSS with 5% FBS and 120 μl of EDTA) and incubated at 37°C with agitation (250 rpm) for 30 minutes. The tissues were filtered using a 100-μm cell strainer and washed with HBSS. Next, the tissues were transferred to 10 ml of digestion buffer (HBSS, 5% FBS, and 50 mg collagenase/Dispase [Roche, Cat# 11097113001, Germany]) and incubated at 37°C with agitation (250 rpm) for another 30 minutes. Following this, the cells were filtered through a 40-μm cell strainer. Finally, the cells were adjusted to 1 million per group and stained for flow cytometry analysis.

### Cytokine gene expression analysis

Ten milligrams of distal colon tissue were collected and homogenized with 1-mm zirconia beads in a Bead-Beater (Fisher Scientific). RNA was extracted using the RNeasy kit (Qiagen), and DNA contaminants were eliminated through DNase digestion (Invitrogen). Approximately 1 μg of RNA was utilized for cDNA synthesis (iSCRIPT; Bio-Rad). Specific primers for pro-inflammatory cytokines (supplemental Table-1) were employed in reverse transcription-PCR (RT-PCR) analysis, conducted using a 7500 Real-Time PCR system (Applied Biosystems). Fold induction was calculated using the threshold cycle (ΔΔCT) method.

### Flow cytometry analysis

Flow cytometry analysis of leukocyte subpopulations was performed as previously described ^9^. In brief, cell suspensions of colonic tissues were initially blocked for 15 minutes with the anti-Fc receptor CD16/CD32 blocking antibody (clone 2.4D2) (BD Pharmingen) at 4°C, followed by incubation with antibodies to CD326, Ly-6G, CD3, panCD45, CD45R0, CD45RA, CD45RB, CD19, CD25, CD69, CD11B, and F4/80. Aqua stain (Life Technologies) was added to identify viable cells. The analysis was conducted on a Becton Dickinson LSR Fortessa. Data were analyzed using FCS Express version 6.0 (De Novo). The gating strategy is illustrated in supplemental Figure 2.

### Statistical analysis

All statistical analyses were performed using GraphPad Prism software. To identify significant differences (∗p < 0.05), parametric data including weight loss, colonization, cytokine expression, and leukocyte infiltration were assessed with a two-way ANOVA followed by Tukey’s multiple comparison test or an unpaired Student’s t-test. For survival curves, the statistics were determined using the log-rank (Mantel-Cox) test with a P value of 0.05.

## Acknowledgments

The study was funded by the UVA Department of Pediatrics and by the National Institute of Allergy and Infectious Diseases (P01AI125181 to F-RP).

## Authorship

Y.L., J.L. A-L., and L.L. conducted the experiments, analyzed the data, and compiled the results. W.G. and G.Z. participated in the experimental design and contributed to scientific discussions. A.E.S., J.P.N., and F.R-P. provided the cell lines, strains, and antibodies and assisted in the experimental design. A.E.S. and F.R-P. wrote the manuscript, conceived the project, and supervised its execution. All authors contributed to the discussions.

## REFERENCES

1. Ruiz-Perez F, Nataro JP. Bacterial serine proteases secreted by the autotransporter pathway: classification, specificity, and role in virulence. Cell Mol Life Sci. 2014;71(5):745–70. doi: 10.1007/s00018-013-1355-8. PubMed PMID: 23689588; PMCID: 3871983.

2. Navarro-Garcia F. Serine proteases autotransporter of Enterobacteriaceae: Structures, subdomains, motifs, functions, and targets. Mol Microbiol. 2023;120(2):178–93. Epub 20230701. doi: 10.1111/mmi.15116. PubMed PMID: 37392318.

3. Debande L, Sabbah A, Kuhn L, Ngondo RP, Maucotel J, Valente-Barroso M, Andre AC, Roche B, Laborde M, Cantalapiedra-Mateo MV, Thahouly T, Milinski A, Bianchetti L, Allmang C, Frugier M, Marteyn BS. SPATEs promote the survival of Shigella to the plasma complement system upon local hemorrhage and bacteremia. Proc Natl Acad Sci U S A. 2024;121(45):e2319951121. Epub 20241030. doi: 10.1073/pnas.2319951121. PubMed PMID: 39475654; PMCID: PMC11551430.

4. Ruiz-Perez F, Wahid R, Faherty CS, Kolappaswamy K, Rodriguez L, Santiago A, Murphy E, Cross A, Sztein MB, Nataro JP. Serine protease autotransporters from Shigella flexneri and pathogenic Escherichia coli target a broad range of leukocyte glycoproteins. Proc Natl Acad Sci U S A. 2011;108(31):12881–6. Epub 2011/07/20. doi: 1101006108 [pii] 10.1073/pnas.1101006108. PubMed PMID: 21768350; PMCID: 3150873.

5. Ayala-Lujan JL, Vijayakumar V, Gong M, Smith R, Santiago AE, Ruiz-Perez F. Broad spectrum activity of a lectin-like bacterial serine protease family on human leukocytes. PLoS One. 2014;9(9):e107920. doi: 10.1371/journal.pone.0107920. PubMed PMID: 25251283; PMCID: 4176022.

6. Hermiston ML, Xu Z, Weiss A. CD45: a critical regulator of signaling thresholds in immune cells. Annu Rev Immunol. 2003;21:107–37. doi: 10.1146/annurev.immunol.21.120601.140946. PubMed PMID: 12414720.

7. Klaus SJ, Sidorenko SP, Clark EA. CD45 ligation induces programmed cell death in T and B lymphocytes. J Immunol. 1996;156(8):2743–53. Epub 1996/04/15. PubMed PMID: 8609392.

8. Liu L, Saitz-Rojas W, Smith R, Gonyar L, In JG, Kovbasnjuk O, Zachos NC, Donowitz M, Nataro JP, Ruiz-Perez F. Mucus layer modeling of human colonoids during infection with enteroaggragative E. coli. Sci Rep. 2020;10(1):10533. Epub 2020/07/01. doi: 10.1038/s41598-020-67104-4. PubMed PMID: 32601325; PMCID: PMC7324601.

9. Vijayakumar V, Santiago A, Smith R, Smith M, Robins-Browne RM, Nataro JP, Ruiz-Perez F. Role of class 1 serine protease autotransporter in the pathogenesis of Citrobacter rodentium colitis. Infect Immun. 2014;82(6):2626–36. doi: 10.1128/IAI.01518-13. PubMed PMID: 24711562; PMCID: PMC4019172.

10. Benjelloun-Touimi Z, Sansonetti PJ, Parsot C. SepA, the major extracellular protein of Shigella flexneri: autonomous secretion and involvement in tissue invasion. Mol Microbiol. 1995;17(1):123–35. Epub 1995/07/01. PubMed PMID: 7476198.

11. Koretzky GA, Picus J, Schultz T, Weiss A. Tyrosine phosphatase CD45 is required for T-cell antigen receptor and CD2-mediated activation of a protein tyrosine kinase and interleukin 2 production. Proc Natl Acad Sci U S A. 1991;88(6):2037–41. doi: 10.1073/pnas.88.6.2037. PubMed PMID: 1672451; PMCID: PMC51163.

12. Trowbridge IS, Thomas ML. CD45: an emerging role as a protein tyrosine phosphatase required for lymphocyte activation and development. Annu Rev Immunol. 1994;12:85–116. doi: 10.1146/annurev.iy.12.040194.000505. PubMed PMID: 8011300.

13. Hieronymus T, Blank N, Gruenke M, Winkler S, Haas JP, Kalden JR, Lorenz HM. CD 95-independent mechanisms of IL-2 deprivation-induced apoptosis in activated human lymphocytes. Cell Death Differ. 2000;7(6):538–47. doi: 10.1038/sj.cdd.4400684. PubMed PMID: 10822277.

14. Lenardo MJ. Interleukin-2 programs mouse alpha beta T lymphocytes for apoptosis. Nature. 1991;353(6347):858–61. doi: 10.1038/353858a0. PubMed PMID: 1944559.

15. Wang R, Rogers AM, Rush BJ, Russell JH. Induction of sensitivity to activation-induced death in primary CD4+ cells: a role for interleukin-2 in the negative regulation of responses by mature CD4+ T cells. Eur J Immunol. 1996;26(9):2263–70. doi: 10.1002/eji.1830260944. PubMed PMID: 8814276.

16. Suzuki H, Kundig TM, Furlonger C, Wakeham A, Timms E, Matsuyama T, Schmits R, Simard JJ, Ohashi PS, Griesser H, et al. Deregulated T cell activation and autoimmunity in mice lacking interleukin-2 receptor beta. Science. 1995;268(5216):1472–6. doi: 10.1126/science.7770771. PubMed PMID: 7770771.

17. Van Parijs L, Biuckians A, Ibragimov A, Alt FW, Willerford DM, Abbas AK. Functional responses and apoptosis of CD25 (IL-2R alpha)-deficient T cells expressing a transgenic antigen receptor. J Immunol. 1997;158(8):3738-45. PubMed PMID: 9103438.

18. Crepin VF, Collins JW, Habibzay M, Frankel G. Citrobacter rodentium mouse model of bacterial infection. Nat Protoc. 2016;11(10):1851–76. doi: 10.1038/nprot.2016.100. PubMed PMID: 27606775.

19. Bhullar K, Zarepour M, Yu H, Yang H, Croxen M, Stahl M, Finlay BB, Turvey SE, Vallance BA. The Serine Protease Autotransporter Pic Modulates Citrobacter rodentium Pathogenesis and Its Innate Recognition by the Host. Infect Immun. 2015;83(7):2636–50. doi: 10.1128/IAI.00025-15. PubMed PMID: 25895966; PMCID: PMC4468532.

20. Chui D, Ong CJ, Johnson P, Teh HS, Marth JD. Specific CD45 isoforms differentially regulate T cell receptor signaling. EMBO J. 1994;13(4):798-807. PubMed PMID: 7509278; PMCID: PMC394878.

21. Dawes R, Petrova S, Liu Z, Wraith D, Beverley PC, Tchilian EZ. Combinations of CD45 isoforms are crucial for immune function and disease. J Immunol. 2006;176(6):3417-25. PubMed PMID: 16517710; PMCID: PMC2619577.

22. Xu Z, Weiss A. Negative regulation of CD45 by differential homodimerization of the alternatively spliced isoforms. Nat Immunol. 2002;3(8):764–71. doi: 10.1038/ni822. PubMed PMID: 12134145.

23. Earl LA, Baum LG. CD45 glycosylation controls T-cell life and death. Immunol Cell Biol. 2008;86(7):608–15. Epub 20080708. doi: 10.1038/icb.2008.46. PubMed PMID: 18607388.

24. Ohta T, Kitamura K, Maizel AL, Takeda A. Alterations in CD45 glycosylation pattern accompanying different cell proliferation states. Biochem Biophys Res Commun. 1994;200(3):1283–9. doi: 10.1006/bbrc.1994.1590. PubMed PMID: 8185577.

25. Liu Z, Dawes R, Petrova S, Beverley PC, Tchilian EZ. CD45 regulates apoptosis in peripheral T lymphocytes. Int Immunol. 2006;18(6):959–66. Epub 20060418. doi: 10.1093/intimm/dxl032. PubMed PMID: 16621865.

26. Perillo NL, Pace KE, Seilhamer JJ, Baum LG. Apoptosis of T cells mediated by galectin-1. Nature. 1995;378(6558):736–9. doi: 10.1038/378736a0. PubMed PMID: 7501023.

27. Xue J, Gao X, Fu C, Cong Z, Jiang H, Wang W, Chen T, Wei Q, Qin C. Regulation of galectin-3-induced apoptosis of Jurkat cells by both O-glycans and N-glycans on CD45. FEBS Lett. 2013;587(24):3986–94. Epub 20131105. doi: 10.1016/j.febslet.2013.10.034. PubMed PMID: 24211831.

